# Self-supervised Component Segmentation To Improve Object Detection and Classification For Bumblebee Identification

**DOI:** 10.1101/2025.03.12.642757

**Authors:** Jahid Chowdhury Choton, Venkat Margapuri, Ivan Grijalva, Brian J. Spiesman, William H. Hsu

## Abstract

The performance of computer vision models for object detection and classification is heavily influenced by the number of classes and quality of input images, particularly in biological applications such as species-level identification of bumblebees. Bee identification is time-consuming, costly, and requires specialized taxonomic training. Different deep learning based computer vision models have been proven to overcome this methodological bottleneck through automated identification of bee species from captured images. However, accurate identification of bee species in images containing multiple objects of various classes poses significant challenges due to ambiguity, poor image quality, and noisy backgrounds. Existing pipelines (baselines) primarily rely on object detection to crop bees from images and classify the species for each cropped instance. This approach is limited by the inclusion of noisy backgrounds, low resolution, and poor image quality. To address these limitations, we propose an enhanced pipeline that integrates object detection with segmentation to generate body masks for bees and remove background noise. This process is complemented by a classification model that identifies the top *k* species for each masked image. The proposed methodology significantly improves both detection and classification performance in most cases, demonstrating its potential to advance automated identification of bee species in complex image datasets. For the cases where the baselines performed much better, we investigated using a state-of-the-art explainable AI model (Grad-CAM) to explain the reason.

## Introduction

Many pollinators, including some bumble bees (i.e., bees in the genus *Bombus*) are experiencing a global decline [1–5]. Bee identification is critical for understanding and preserving biodiversity, monitoring pollination networks, and supporting agricultural ecosystems [3, 4]. The decline of bees poses significant threats to food security and ecological balance. Traditional methods of bee identification, which rely on expert taxonomists and manual inspection, are time-consuming, labor-intensive, and can be prone to human error [5, 6]. The advent of deep learning and computer vision has helped to revolutionize this field by enabling automated, accurate, and scalable identification of bee species. Leveraging advanced models for object detection, segmentation, and classification, these technologies can analyze large datasets of bee images, even in challenging conditions such as cluttered backgrounds or poor image quality. But current classification models for bee species identification have much room for improvement. By leveraging hierarchical approaches, computer vision approaches hold immense potential to enhance species-level identification, facilitate real-time monitoring, and contribute to data-driven conservation efforts on a global scale.

Preparing datasets for bees with multiple objects requires manual annotation of each instance on each image. Moreover, training state-of-the-art computer vision models such as YOLOv9 [7] for object detection and classification can be time-consuming and resource-intensive, particularly for datasets with a large number of classes (*>*100). To solve this problem, Spiesman *et al*. [6, 8] developed a pipeline where the object detection and classification are conducted separately in two different models, reducing the time and resource overheads. At first, YOLOv9 model is trained on a single class label (bee) for object detection. Next, each instance of the bees is detected and cropped out from the image so that there is only one bee in each cropped image. The cropped images are used to train a classification model (EfficientNetV2 [9]) which is much smaller and faster than the object detection model. The trained pipeline is deployed as a web application called Beemachine (https://beemachine.ai/) which can output the highest probability of the top three species of a bee from a given set of images. However, cropped images can have low resolutions, noisy backgrounds, and poor qualities. Therefore, further improvement and pre-processing of these cropped images can help to improve the accuracy of the classification model.

In this research, we propose a new pipeline for identifying bee species, which includes improving the object detection and classification of Bumblebees through background removal performed by mask segmentation. We also improve object detection by combining the supervised model YOLOv9 [7] trained on annotated datasets, and the zero-shot model GroundingDINO [10]) fine-tuned on the same dataset. First, we crop each bee instance from the input image using the improved object detection model. Next, we fine-tune Segment-Anything-2 (SAM-2) [11], a state-of-the-art segmentation model for generating a mask segment of the full bee body in each cropped image, and remove the background by replacing the pixels outside the bee with zeros (black). Finally, we train the classification model EfficientNetV2 [9] to detect the probability of top *k* species for each segmented image. Here, *k* is the number of species that have more than 5% probability of being present in an image. The baseline (blue arrows) and the proposed pipeline (orange arrows) are illustrated in Fig. 1. Several experiments are performed in each stage of the proposed pipeline, and compared with the existing pipeline. Furthermore, we investigate the cases where the baseline performs better by checking the feature map of the classification model using an explainable AI tool named Grad-CAM [12]. Through the experiments, we showed that our proposed pipeline that perform multi-stage detection by mask segmentation offer better performance compared to existing independent models.

**Fig. 1:**
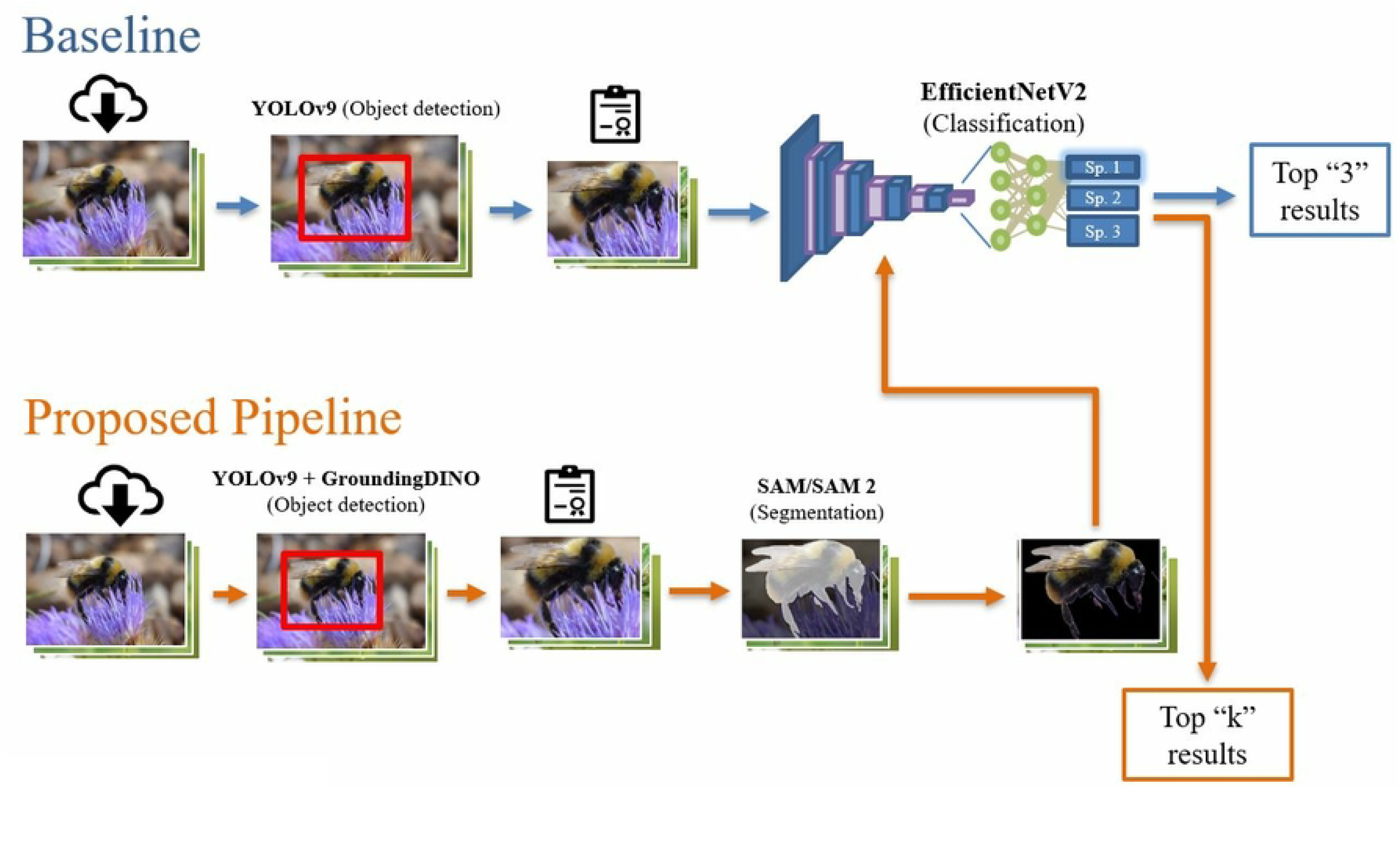
Overview of the Crop-to-Mask pipeline for bee identification

### Related work

Computer vision and deep learning algorithms have revolutionized species-level identification, offering promising solutions for automated species recognition and monitoring [6, 13–16]. Convolutional neural networks (CNNs) have emerged as the most effective approach for automated classification and detection using image data [9, 16–20]. But in traditional classification datasets, (ImageNet [21], CIFAR100 [22], etc.) each image only belongs to a single class. An image containing various objects belonging to different classes can be significantly more challenging [15, 17, 23]. Therefore, object detection is a fundamental task in computer vision that aims to identify and localize objects of interest within an image or video frame. Many efficient object detection models have been developed in the last decade such as YOLO [23], Region-based CNN [24], Fast R-CNN [25], Faster R-CNN [26], and so on. However, these models usually require large datasets and manual annotation of each object for training. In the past few years, few-shot learning [27–33] and zero-shot learning [10, 34–43] have emerged as popular alternatives to reduce the labor-intensive and time-consuming task to make large datasets. Few-shot learning is a framework in which a model learns to make accurate predictions by training on a few labeled examples. On the other hand, zero-shot learning is a framework that allows models to recognize new classes without being trained on examples of those classes. In most object detection models, the objects are usually detected within bounding boxes covering that object. However, the shapes of the objects may be irregular, and the bounding boxes may contain noisy backgrounds that are not relevant to the original object. Therefore, image segmentation is a task that focuses on dividing an image into regions belonging to different classes at the pixel level, resulting in irregular shapes [44–49]. Recently, the Segment Anything Model (SAM) [50] and its successor SAM 2 [11] have been widely used for segmentation in domains such as medical imaging [48, 51–53], agricultural imaging [49, 54–56], and so on. SAM and SAM 2 models can take the bounding boxes provided by object detection models as inputs, and create a segmentation mask that contains the object without any background. Our idea is to combine two state-of-the-art object detection models (Yolov9 [7] and GroundingDINO [10]) for generating the bounding boxes taken as inputs by the state-of-the-art segmentation model SAM 2 [11] to generate masks of bees. The Grounded SAM and Grounded SAM 2 [57] are two models that can generate bee body masks using the zero-shot approach. But since we already have a trained supervised model developed by Spiesman *et al*. [6, 8] for object detection, we want to leverage its advantage by combining it with the zero-shot model GroundingDINO [10]. There have been many efficient classification models based on CNNs that are smaller and faster to train and do not have the bottleneck of object detection [9, 18, 19, 58]. Among others, EfficientNetV2 [9] achieves top state-of-the-art performance on the ImageNet [21] and CIFAR100 [22] datasets that are widely used as a benchmark for testing the performance of classification models. In this research, we leverage the benefits of combining state-of-the-art computer vision models sequentially by dividing the species detection task into a sequence of three subtasks: object detection, segmentation, and classification. We use YOLOv9 [7] and GroundingDINO [10] for object detection, Segment Anything 2 (SAM 2) for segmentation, and EfficientNetV2 [9] for classification. This novel species detection pipeline will only focus on features that are present inside the bee body, and improve the accuracy of the bee species detection.

## Materials and Methods

The image dataset used for the experiment is the updated version of the bumblebee dataset described by Spiesman et al. [6]. It contains 195,185 images belonging to 166 different *Bombus* species. Each original image has at least 200×200 pixels. Different subsets of this dataset are used for training and fine-tuning different computer vision models. The first subset comprises 32,344 high-quality images with bounding box annotations for object detection. YOLOv9 [7] is trained and GroundingDINO [10] is fine-tuned on this subset where 80% images are used for training, 10% images for validation and 10% for testing. Images are resized to 225×225 pixels and pre-processed by standardizing pixel values for each RGB channel between 0 and 255. Several image augmentation techniques are used to reduce overfitting and increase the generalizability of the model: rotation (≤ 100°), shifting (≤ 25%), flipping, random cropping, and random channel inversion. The YOLOv9 model is trained for 100 epochs with an initial learning rate of 0.01, batch size of 64, momentum of 0.9, and a decay of 0.0005 whenever the validation loss reached a plateau. For the optimizaiton, the *AdamW* [59] algorithm is used.

Next, we generated a small subset of 5,478 images with mask annotations in the VGG format [60]. This dataset contains the precise full-body segmentations of the bees. We generated binary mask images from the VGG annotations suitable for fine-tuning the segmentation models SAM [50] and SAM 2 [11]. The number of training, validation, and testing images was split in a ratio of 80%:10%:10%. Other than the number of epochs (which is 200), we use the same hyperparameters as the object detection model for fine-tuning SAM 2.

In the baseline approach, the classification model (EfficientNetV2) was trained on the full dataset with a splitting ratio of 85%:10%:5% for training, validation, and testing respectively. In our proposed pipeline, we used the same splitting ratio. But the classification model takes a masked image (no background) as input, as shown in Fig. 1. Therefore, we generated the masked image dataset from the original dataset by object detection and segmentation as illustrated in Fig. 2. For object detection, we maximized recall by combining YOLOv9 and GroundingDINO models to generate box annotations. These box annotations were taken as inputs to generate the segmentation masks using SAM and SAM 2, and the backgrounds were removed for each image. The following sections describe the details of the object detection and segmentation steps. The object detection models couldn’t recognize any bee for a very small number of images (222) within the full dataset. Therefore, the number of images in our masked dataset is: 1,95,185 - 222 = 1,94,963. The masked dataset was split in the same ratio as before, and the details are described in Table 1. We trained the classification models for 200 epochs using the *Adam* [61] optimizer with an initial learning rate of 0.01, batch size of 64, weight decay of 0.001, momentum of 0.9, dropout of 0.2, and the same image augmentation techniques used in YOLOv9. The classification models are used to detect the probability of top *k* species on an input image, where *k* is the number of species that have more than 5% probability of being present in the image.

**Table 1.** Datasets for different CV models.

| Task | Models | Training | Validation | Testing | Total |
| --- | --- | --- | --- | --- | --- |
| Object Detection | YOLOv9, GroundingDINO | 25,875 | 3,235 | 3,234 | 32,344 |
| Segmentation | SAM, SAM 2 | 3,314 | 1,082 | 1,082 | 5,478 |
| Classification (Existing) | EfficientNetV2 | 1,65,907 | 19,518 | 9,760 | 1,95,185 |
| Classification (Proposed) | EfficientNetV2 | 1,65,704 | 19,518 | 9,760 | 1,94,982 |

**Fig. 2:**
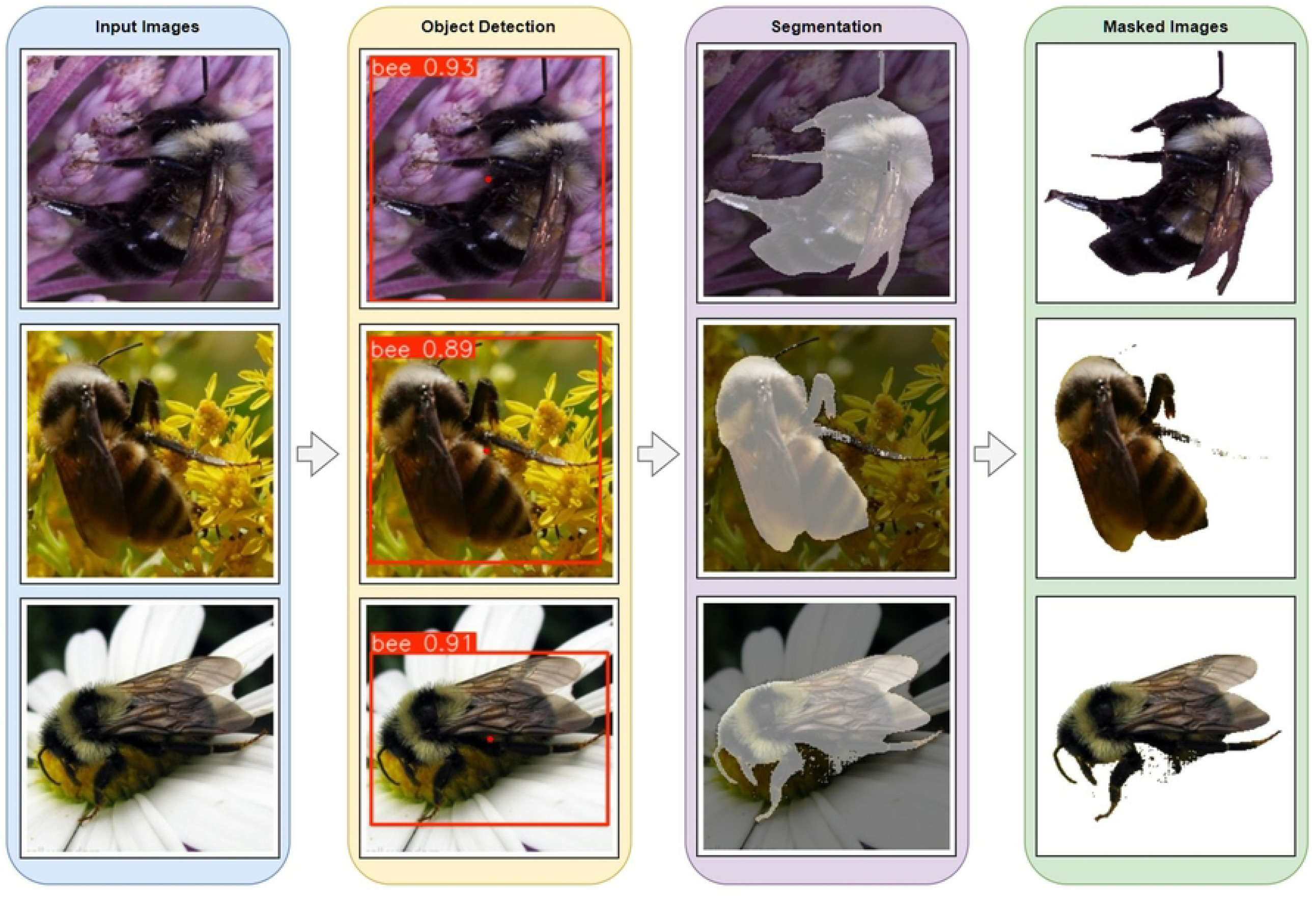
Generating masked dataset from the original dataset. Each image is passed through the object detection and the segmentation models to remove the backgrounds.

### Object detection

Object detection is a fundamental task in computer vision that involves identifying and localizing objects within an image. YOLOv9 [7], a successor to YOLOv8, is a state-of-the-art supervised object detection model which is well-known for its high speed and accuracy. It requires labeled datasets where bounding boxes and class labels are manually annotated. YOLOv9 incorporates advancements in architectural design, such as the concept of programmable gradient information (PGI), improved backbone networks, dynamic anchor-free mechanisms, and attention-based feature aggregation. Even though YOLOv10 has already been released, its performance is worse than YOLOv7/v9 on custom datasets [62]. Therefore, we selected and trained YOLOv9 on a subset of the full bee dataset with box annotations (32,344 images) having a single class label (bee) as mentioned before.

The term zero-shot refers to a machine learning approach where a model can perform a task or make predictions for classes, categories, or tasks it has never explicitly seen or been trained on before. This capability is achieved by leveraging generalization and prior knowledge from pretraining on a large and diverse dataset. In our case, we use the text prompt “bee” as the input of GroundingDINO to detect all “bee” instances in a given image, and then compare the output with the YOLOv9 model. In the baseline approach, the YOLOv9 model is trained to detect all the bees in a given input image, and each detection of the bee is cropped to create images where there is only one bee per image. As mentioned in [8], this model achieves 99.5% accuracy in bee detection for pinned bee specimens. However, if the variations in the images increase (images coming from new species), we may need to retrain the model even though the model is trained on a single class, and the accuracy may go down. Moreover, we will need to make more box annotations for new images. This can be labor-intensive and time-consuming. Therefore, we propose to use a state-of-the-art open-set object detection model named GroundingDINO [10]. GroundingDINO is a zero-shot model that uses vision-language pretraining. It can generalize to novel objects and tasks without requiring explicit retraining. The model leverages transformer-based architectures and grounding mechanisms to align textual descriptions with visual features. Grounding DINO was trained on a large and diverse dataset of approximately 1.8 million images. These images were sourced from various publicly available datasets and include both object detection and grounding annotations. The latest API of Grounding DINO (version 1.6) has achieved the best performance in MSCOCO [63] and LVIS-minimal [64] datasets.

To compare the performance of YOLOv9 and GroundingDINO, we assess both localization and classification accuracy using the following metrics: intersection-over-union (IoU), precision, recall, and F1-score. The intersection-over-union or IoU is the ratio between the overlap of predicted and ground truth bounding boxes and the union of these two bounding boxes. Precision measures the accuracy of positive predictions. It is calculated as the number of true positives divided by the total number of positive predictions or *TruePositives/*(*TruePositives* + *FalsePositives*). Recall measures the ability to find all ground-truth objects and is calculated as *TruePositives/*(*TruePositives* + *FalseNegatives*). The F1-score is the harmonic mean between precision and recall and is calculated as (2 × *precision* × *recall*)*/*(*precision* + *recall*). The number of detections (recall) is maximized by sequentially running inference on YOLOv9 and GroundingDINO. In the majority of images, both models detect bees accurately. In cases where both models detect bee(s), the detections made by YOLOv9 are considered for further analysis. The images in which either model fails to detect bees are ignored and not analyzed further.

### Segmentation

Segmentation is a computer vision task aimed at dividing an image into meaningful regions or segments. Each segment represents a specific object, part of an object, or a region of interest within the image. Segmentation is used to achieve a more detailed understanding of the visual content. Full body segmentation of bees gives us the exact pixels of a bee body, and does not have background pixels present in a bounding box annotation. The Segment Anything Model (SAM) [50] and its advanced version, SAM 2 [11], are cutting-edge segmentation models designed to address a wide range of computer vision tasks. SAM, developed by Meta AI, is a foundation model for segmentation that excels in interactive and prompt-based segmentation tasks. It can generalize across diverse datasets, making it adaptable to various domains without requiring additional fine-tuning. SAM 2 builds on its predecessor by introducing improved efficiency, better handling of fine-grained details, and support for multimodal inputs, enabling enhanced segmentation performance even in complex scenarios. Both models are valuable for applications such as medical imaging, object recognition, and environmental monitoring, where precise and adaptable segmentation is crucial [48, 49, 51–56]. As shown in Fig. 1, we take the cropped bee images as inputs, and remove the background by sending it to the segmentation model (SAM/SAM 2). All pixels outside the bee body are replaced with zero for all channels, resulting in a black background. This process ensures that the classification model only focuses on the pixels inside the bee body while training, and prevents itself from learning the external environment. Furthermore, it makes the size of each cropped image significantly smaller, resulting in faster inference and training speed. Even though SAM and SAM 2 support zero-shot prompting, we fine-tuned both models using a small hand-segmented dataset (5,478 images).

### Classification

Classification is a supervised learning task where the goal is to assign a label or category to input data based on its features. A classification model learns from labeled training data and uses the learned patterns to predict the label for new, unseen data. Convolutional Neural Networks (CNNs) are widely used for image classification because of their ability to automatically and efficiently extract features from images. EfficientNetV2 is a family of convolutional neural networks that have been leveraged for classification tasks in this work, specifically using its small (S), medium (M), and large (L) variants. Compared to traditional convolutional neural networks, EfficientNetV2 achieves better accuracy with fewer parameters, reducing memory requirements. EfficientNetV2 introduces fused-MBConv layers, which enhance the training and inference speed while keeping the network size smaller [9]. The model performs well across various domains, including natural images and fine-grained classification tasks, making it an ideal choice for general-purpose and domain-specific datasets. In the baseline approach, the three variants of EfficientNetV2 were trained on the full dataset for bee identification. However, in our proposed pipeline, we first generate the masked dataset using segmentation and then train these models on it. As shown in Fig. 2, each image in the masked dataset has the pixels of a bee body without any background. The masks have small errors (± 0.02%) that either cut-off the bee body or include the background (compared to the ground truth masks). Even with these errors, we successfully prevent the classification models from learning the features that are not present inside the bee body. We hypothesize and confirm that this results in faster training speed and accurate learning of the desired object.

### Experiments

We performed three computational experiments using our proposed pipeline for the three tasks: object detection, segmentation, and classification. For object detection, the performance between GroundingDINO and YOLOv9 is compared by measuring the precision, recall, F1 score, and intersection-over-union (IoU) score for the confidence thresholds of 0.25, 0.35, 0.5, and 0.75. The experimental results are shown in Table 2 and 3. Although our YOLOv9 model is trained on a smaller dataset, we run inference on the full dataset and measure its performance. Since YOLOv9 is a supervised model, we expected it to perform better than GroundingDINO. However, the F1 score at the confidence threshold of 0.35 is much higher in GroundingDINO. This confirms that zero-shot detection can be used efficiently by tuning the confidence thresholds, eliminating the need for supervised models. For confidence thresholds 0.25 and 0.35, GroundingDINO had a lot of false positives (the number of detections are higher than number of objects), which indicates that we must be careful even when the F1 score is high as the model detects a bounding box around the bee. Both models achieve 100% precision at thresholds 0.5 and 0.75. For IoU scores in Table 3, GroundingDINO performed much better than YOLOv9, resulting in tighter bounding boxes surrounding the bee.

**Table 2.** Precision, Recall, and F1 scores between GroundingDINO and YOLOv9.

| No | Confidence | Total Objects | GroundingDINO |  |  |  |  |  | YOLOv9 |  |  |  |  |  |
| --- | --- | --- | --- | --- | --- | --- | --- | --- | --- | --- | --- | --- | --- | --- |
|  |  |  | Det. | FP | FN | Prec. | Recall | F1 | Det. | FP | FN | Prec. | Recall | F1 |
| 1 | 0.25 | 195185 | 268244 | 73059 | 0 | 0.7859 | 1 | 0.8801 | 188155 | 7 | 7037 | 0.9999 | 0.9652 | 0.9822 |
| 2 | 0.35 | 195185 | 201152 | 5971 | 4 | 0.9711 | 0.9999 | 0.9853 | 186379 | 6 | 8812 | 0.9999 | 0.9568 | 0.9779 |
| 3 | 0.5 | 195185 | 153645 | 0 | 41540 | 1 | 0.8245 | 0.9038 | 183549 | 0 | 11636 | 1 | 0.9437 | 0.9710 |
| 4 | 0.75 | 195185 | 117194 | 0 | 77991 | 1 | 0.7145 | 0.8334 | 173064 | 0 | 22121 | 1 | 0.8982 | 0.9463 |

**Table 3.** IoU scores between GroundingDINO and YOLOv9.

| No | Confidence | GroundingDINO |  |  | YOLOv9 |  |  |
| --- | --- | --- | --- | --- | --- | --- | --- |
|  |  | Mean | Median | SD | Mean | Median | SD |
| 1 | 0.25 | 0.917598835 | 0.955071688 | 0.111446316 | 0.454645524 | 0.48014544 | 0.223305575 |
| 2 | 0.35 | 0.90190396 | 0.956214368 | 0.172988115 | 0.453793329 | 0.48014544 | 0.224396144 |
| 3 | 0.5 | 0.775622191 | 0.949893415 | 0.363159543 | 0.451825563 | 0.47537734 | 0.226227864 |
| 4 | 0.75 | 0.615332197 | 0.934286982 | 0.457005616 | 0.436179473 | 0.46582651 | 0.239792384 |

For segmentation, we compare the performance between SAM and SAM 2 by measuring IoU scores for the same confidence thresholds. The results are shown in Table 4. SAM 2 outperforms SAM for all confidence thresholds. The average IoU scores increased as we increased the confidence threshold and vice versa, as hypothesized. However, after reaching a fixed value, the IoU scores no longer improved (almost the same for confidence thresholds 0.5 and 0.75). Therefore, the confidence threshold is set to 0.5 to detect a mask segmentation that is close to the true segmentation. Still, for many images (∼ 8,000), the SAM 2 model predicted incorrect or bad masks which contain at least one of the following deficiencies: too small, too large, missing body parts, including backgrounds, and multiple contours. All these images have low-resolution and poor quality. Therefore, we removed these noisy images from the masked dataset.

**Table 4.** IoU scores between SAM and SAM 2.

| No | Confidence | SAM |  |  | SAM 2 |  |  |
| --- | --- | --- | --- | --- | --- | --- | --- |
|  |  | Mean | Median | SD | Mean | Median | SD |
| 1 | 0.25 | 0.265276165 | 0.08009828 | 0.294226752 | 0.459894292 | 0.392791846 | 0.433454346 |
| 2 | 0.35 | 0.202662254 | 0.069998741 | 0.30900157 | 0.487528126 | 0.489364843 | 0.442158792 |
| 3 | 0.50 | 0.320429984 | 0.080056845 | 0.343198383 | 0.597424779 | 0.841315032 | 0.392966555 |
| 4 | 0.75 | 0.322724851 | 0.080811955 | 0.34399464 | 0.598611777 | 0.838779136 | 0.39851788 |

For classification, we perform the experiments on the original dataset for the baseline approach, and on the masked dataset for our proposed pipeline. The three variations of the EfficientNetV2 model (EfficientNetV2S, EfficientNetV2M, and EfficientNetV2L) were trained and tested with identical preprocessing, hyperparameters, and data augmentation techniques described in section 2 to ensure a fair comparison. For the baseline approach, EfficientNetV2M outperformed both EfficientNetV2S and EfficientNetV2L as shown in Table 5. This proves that even though EfficientNetV2L has higher accuracy in the ImageNet dataset [21], smaller models can sometimes perform better on custom datasets where the variation between different classes is very small and specific. For the proposed pipeline, EfficientNetV2L outperformed the other two models by a large margin (∼ 11% gain in mAP). Comparing the existing and proposed pipeline, we received 14% and 10% increase in mAP for EfficientNetV2S and EfficientNetV2L respectively, but got nearly 3% less accuracy in EfficientNetV2M. Therefore, further investigation is necessary to explain why the “bee image with background” performed better than the “bee image with no background”. Fig. 3 shows the details.

**Table 5.** Distributive focal loss (DFL) and mean average precision (mAP) for EfficientNetV2S, EfficientNetV2M, and EfficientNetV2L on the baseline and proposed pipeline.

| No | Model | Baseline |  | Proposed Pipeline |  |
| --- | --- | --- | --- | --- | --- |
|  |  | Loss | mAP | Loss | mAP |
| 1 | EfficientNetV2S | 1.5402 | 0.6303 | 0.8146 | 0.7705 |
| 2 | EfficientNetV2M | 0.7763 | 0.7601 | 0.8301 | 0.7387 |
| 3 | EfficientNetV2L | 0.6954 | 0.7425 | 0.5422 | 0.8489 |

**Fig. 3:**
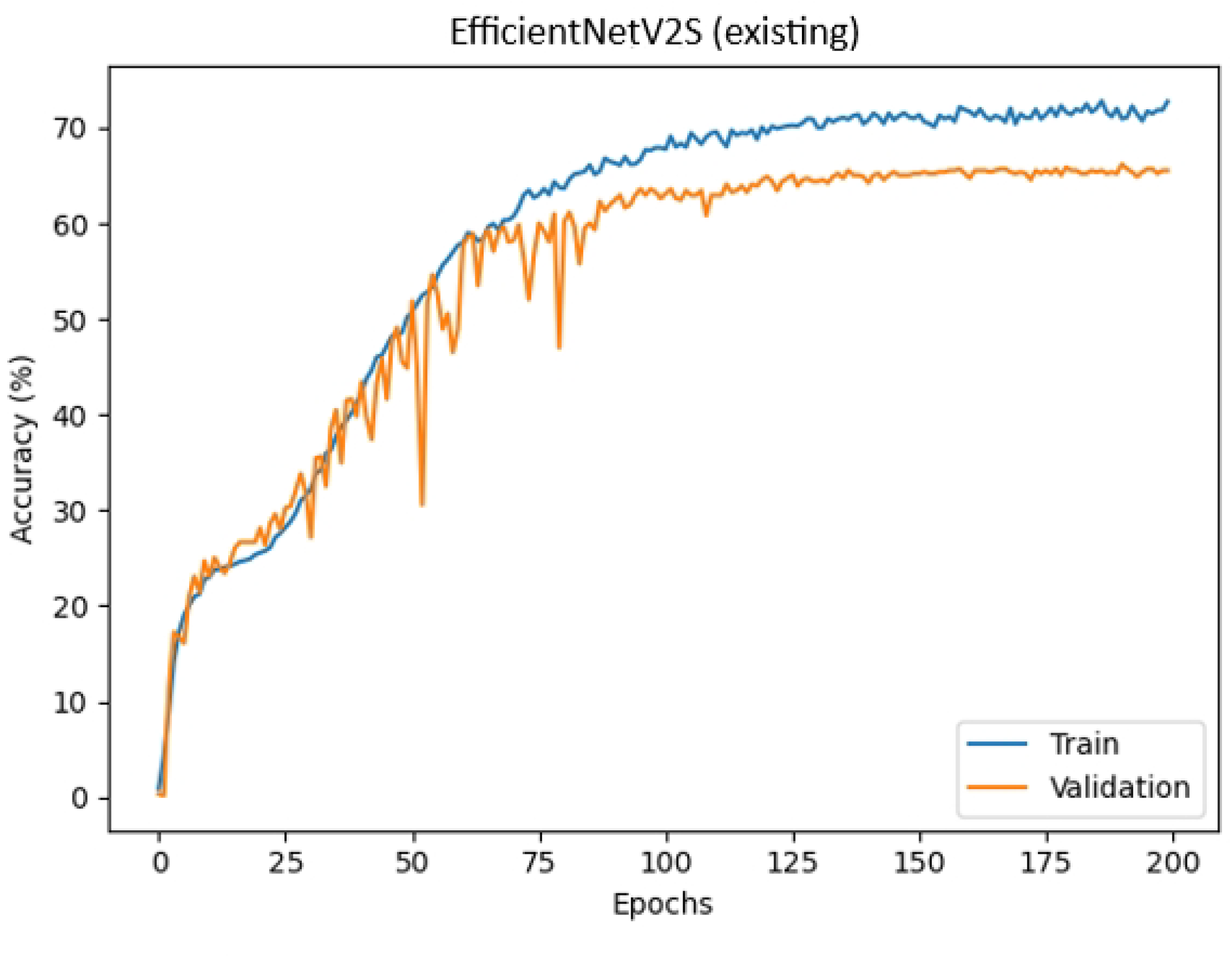
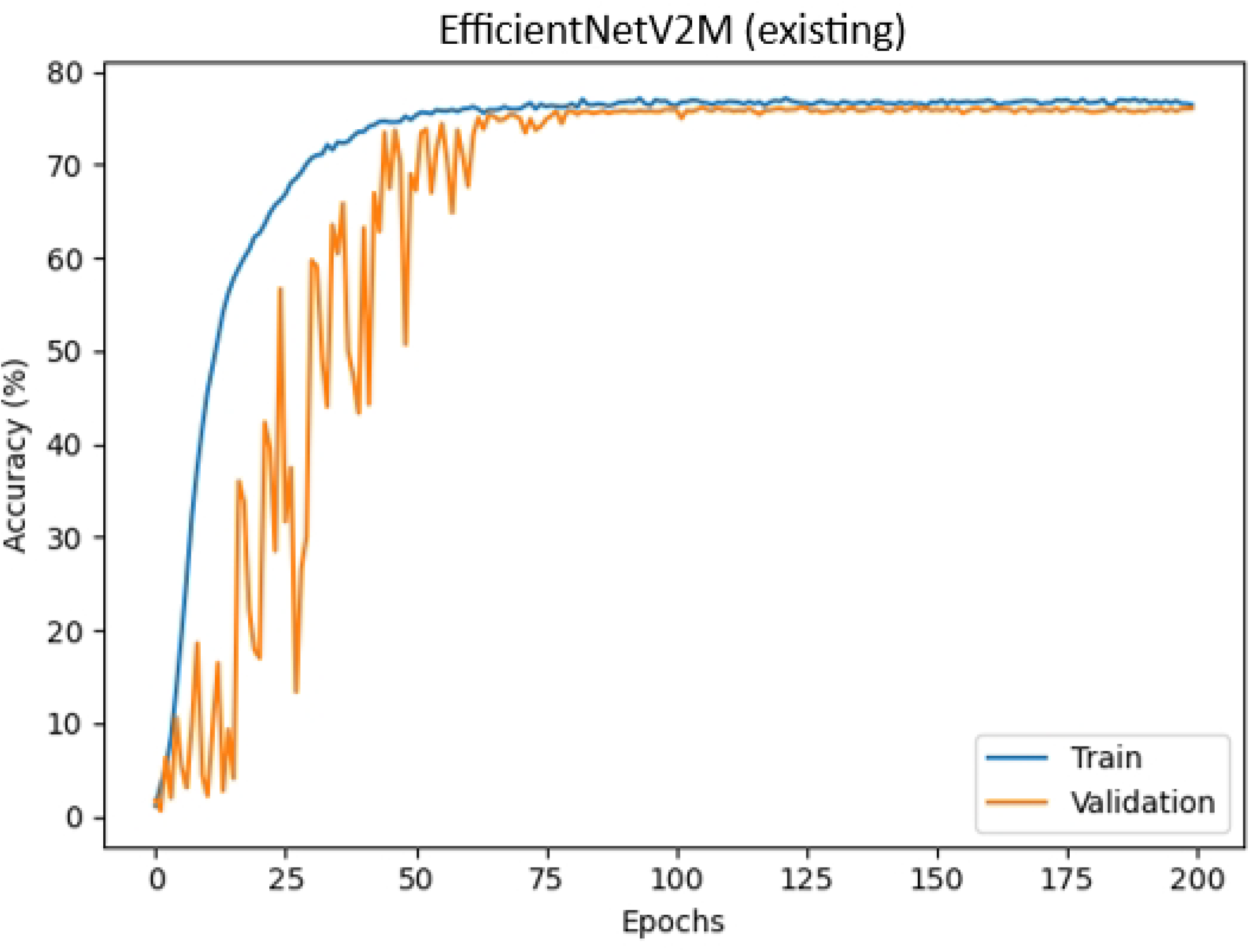
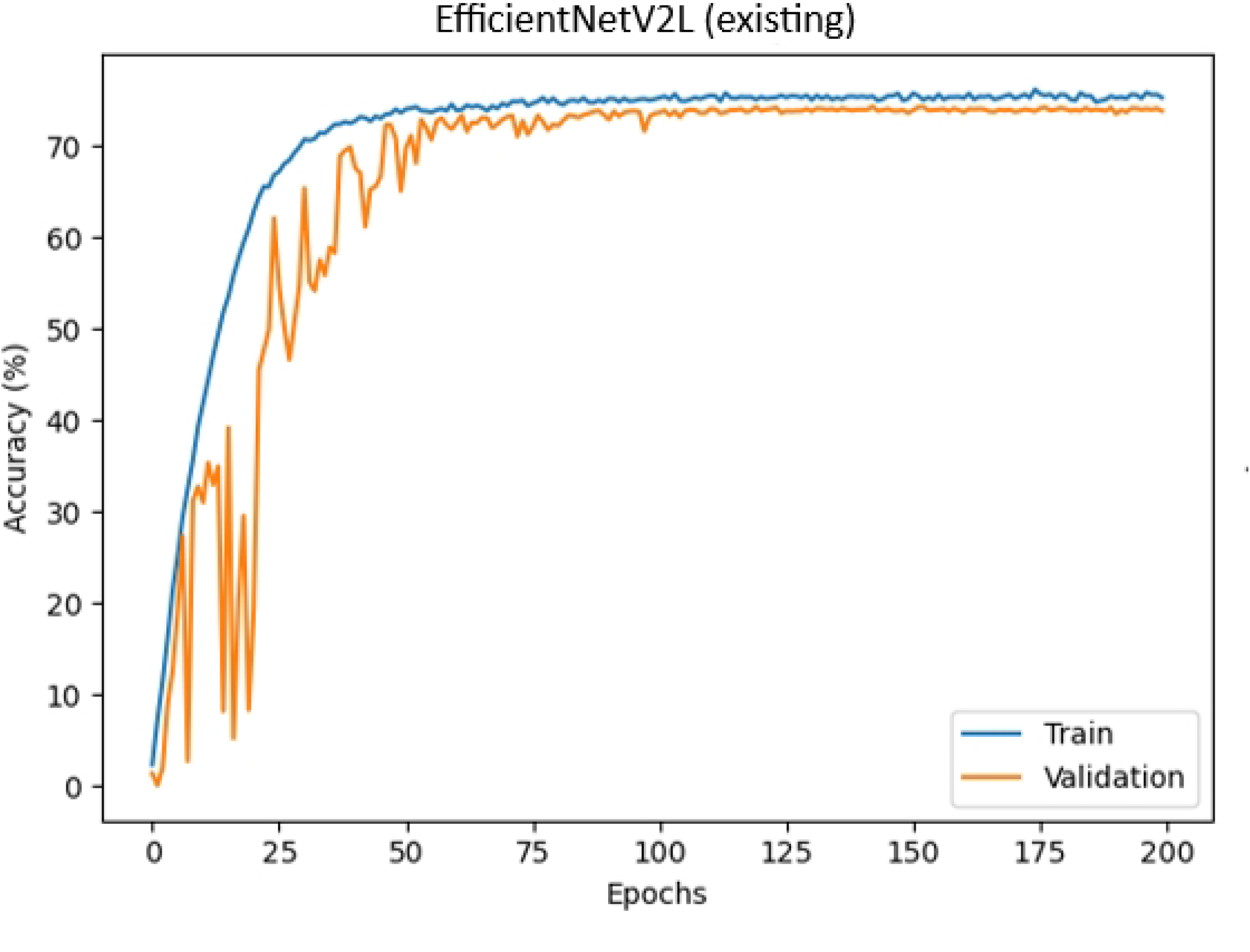
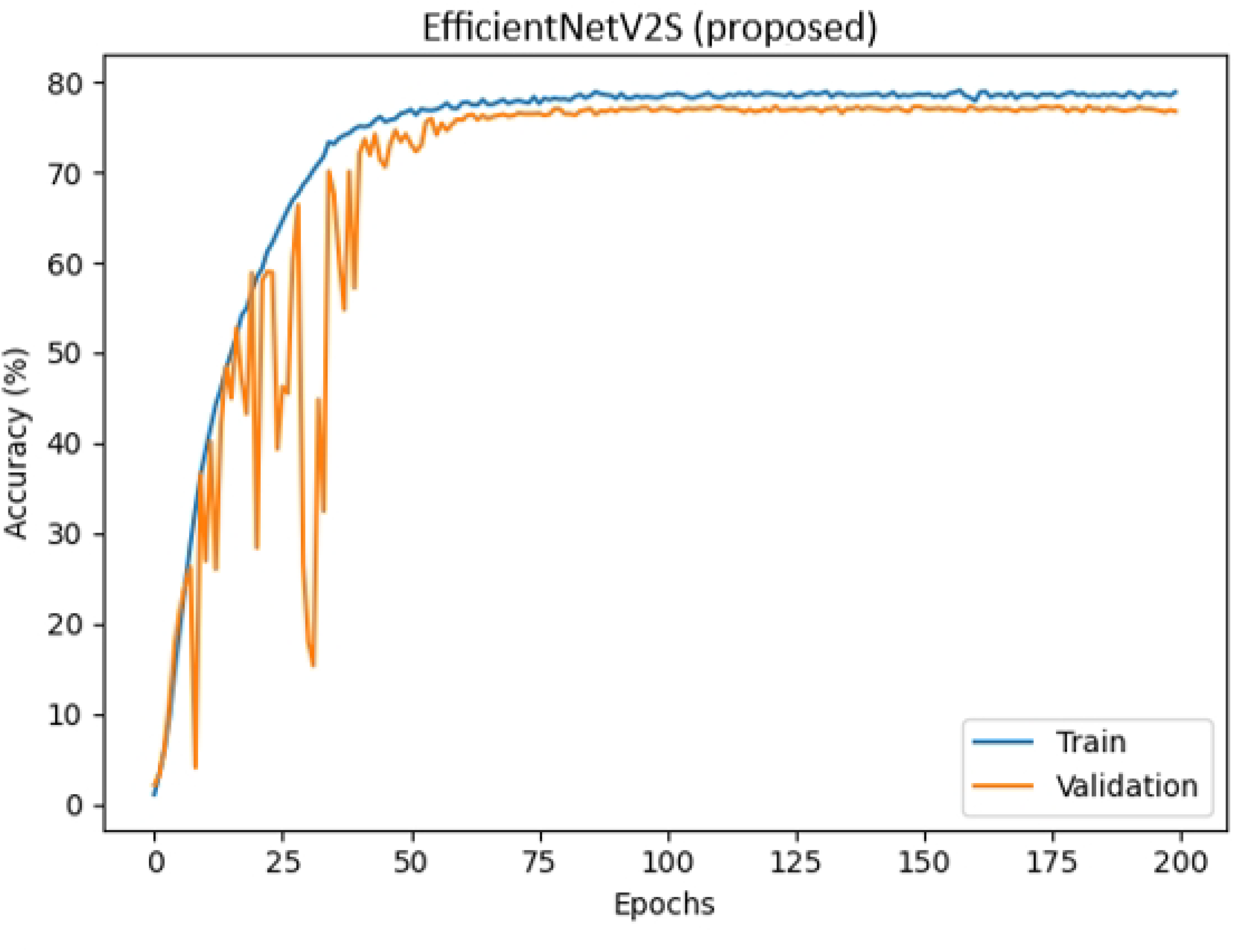
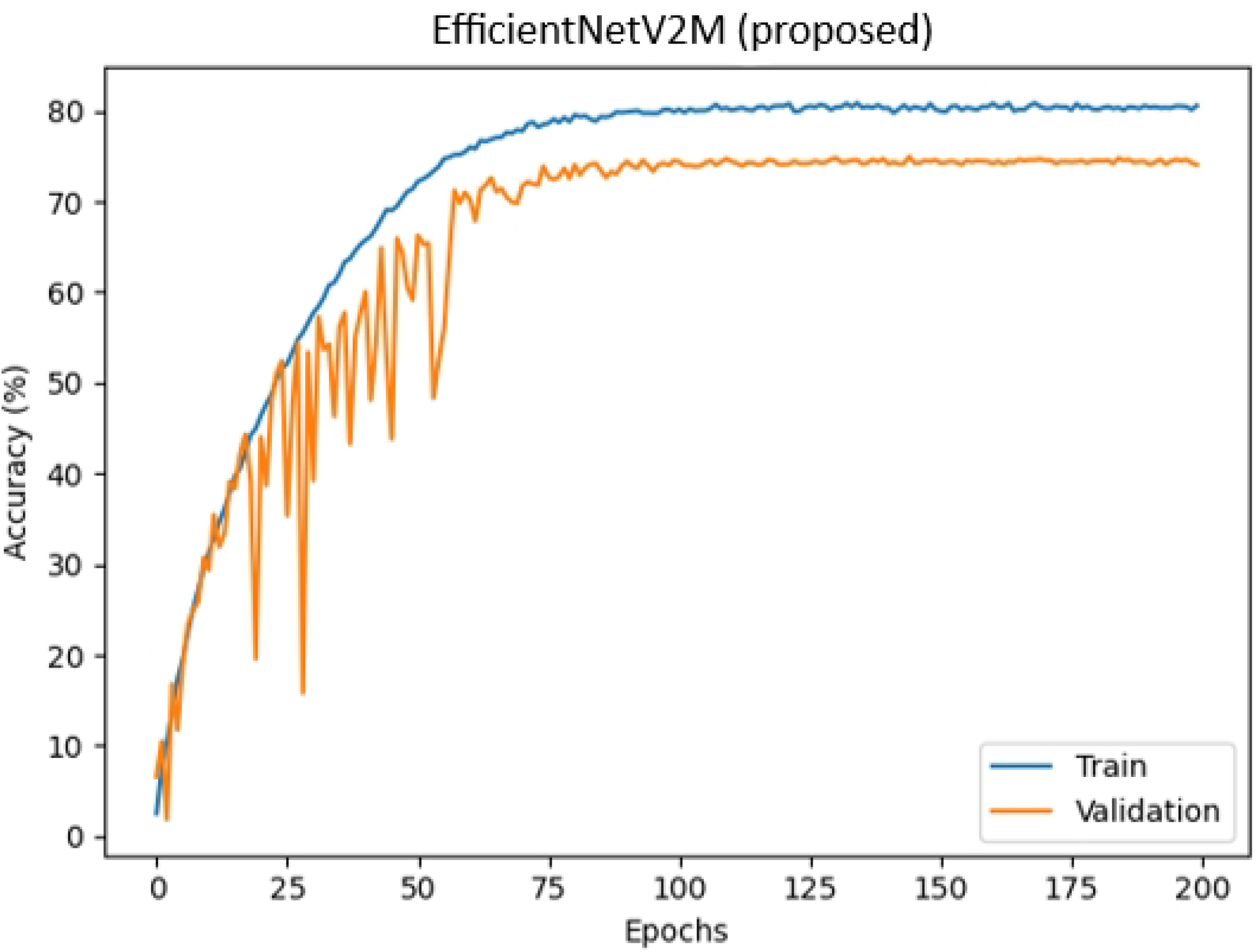
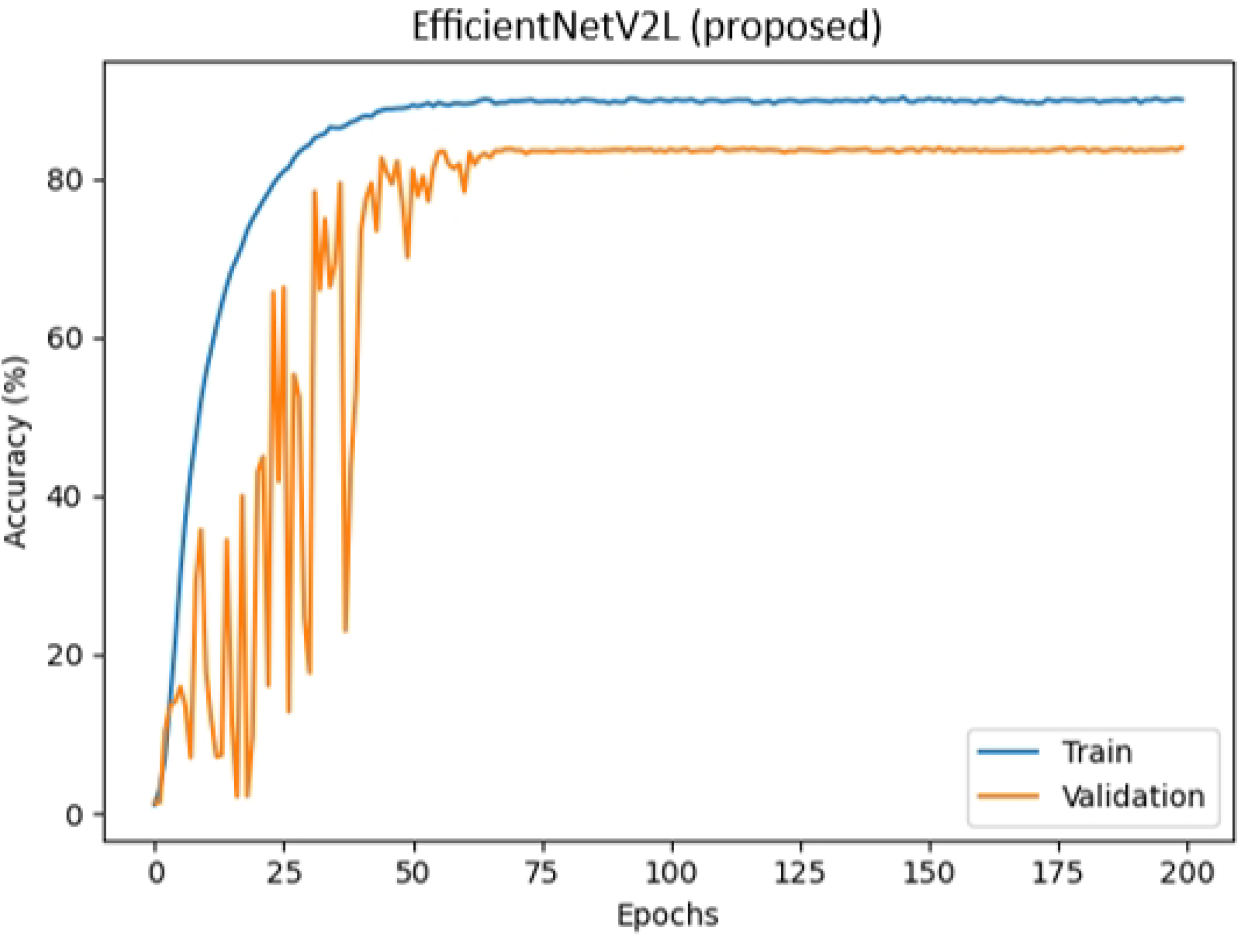
Figure (a), (b), and (c) are the accuracy plots for the existing pipeline, and Figure (d), (e), and (f) are the accuracy plots for the proposed pipeline using EfficientNetV2S, EfficientNetV2M, and EfficientNetV2L classification models respectively.

### Investigation: Grad-CAM

Gradient-weighted Class Activation Mapping (Grad-CAM) [12] is a widely used visualization technique to interpret convolutional neural networks (CNNs) by highlighting regions in the input image that are most influential in a model’s decision-making process. In the context of EfficientNetV2, Grad-CAM can provide valuable insights into the features learned by the network and the areas it focuses on while performing classification tasks. Since the original model (bee image with background) performed better than the proposed model (bee image with no background) in one of the vision models (EfficientNetV2M), it means that the model must be learning something outside of the bee body, and that “something” is also present on the test images, giving it correct prediction for bee species. This assumption is confirmed by the Grad-CAM analysis shown in Figure 4. Here in Fig. 4 (a), the highlighted region is within the bee body, therefore making the correct prediction for the species (*Bombus affinis*). However, in Fig. 4 (b), the highlighted region is in front of the bee head, and irrelevant to species detection. The reason why the CV models can correctly detect the species even when focusing outside the bee, is that many bee species usually visit the same plant/flower. Therefore, focusing on that flower can help in detecting that species. But when the flower is absent from the image, the model may make incorrect predictions as shown in Fig. 4 (b). The correct species is *Bombus affinis*, but the model predicted it as *Bombus appositus*. This investigation provides valuable information on what the model had actually learned.

**Fig. 4:**
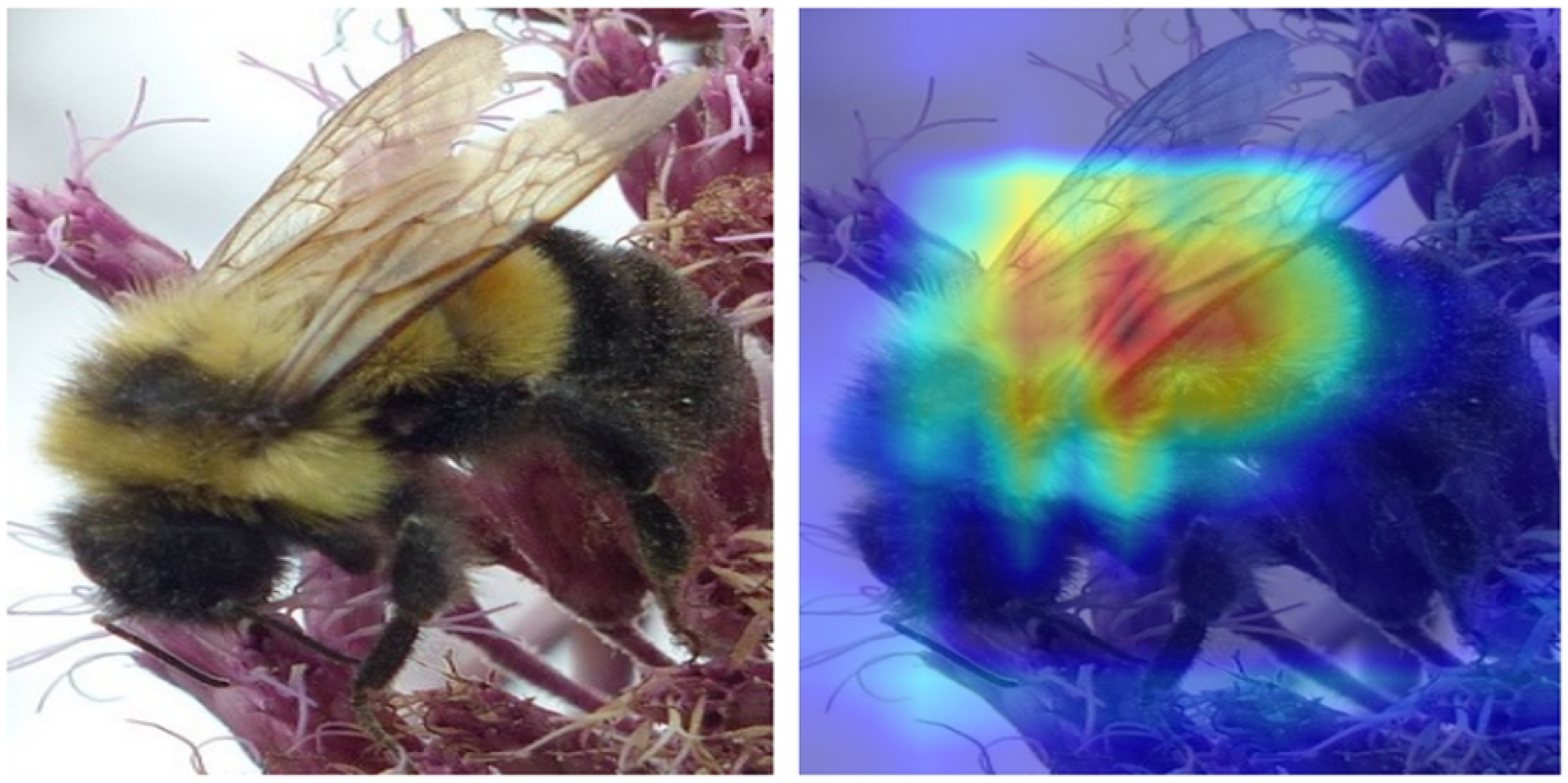
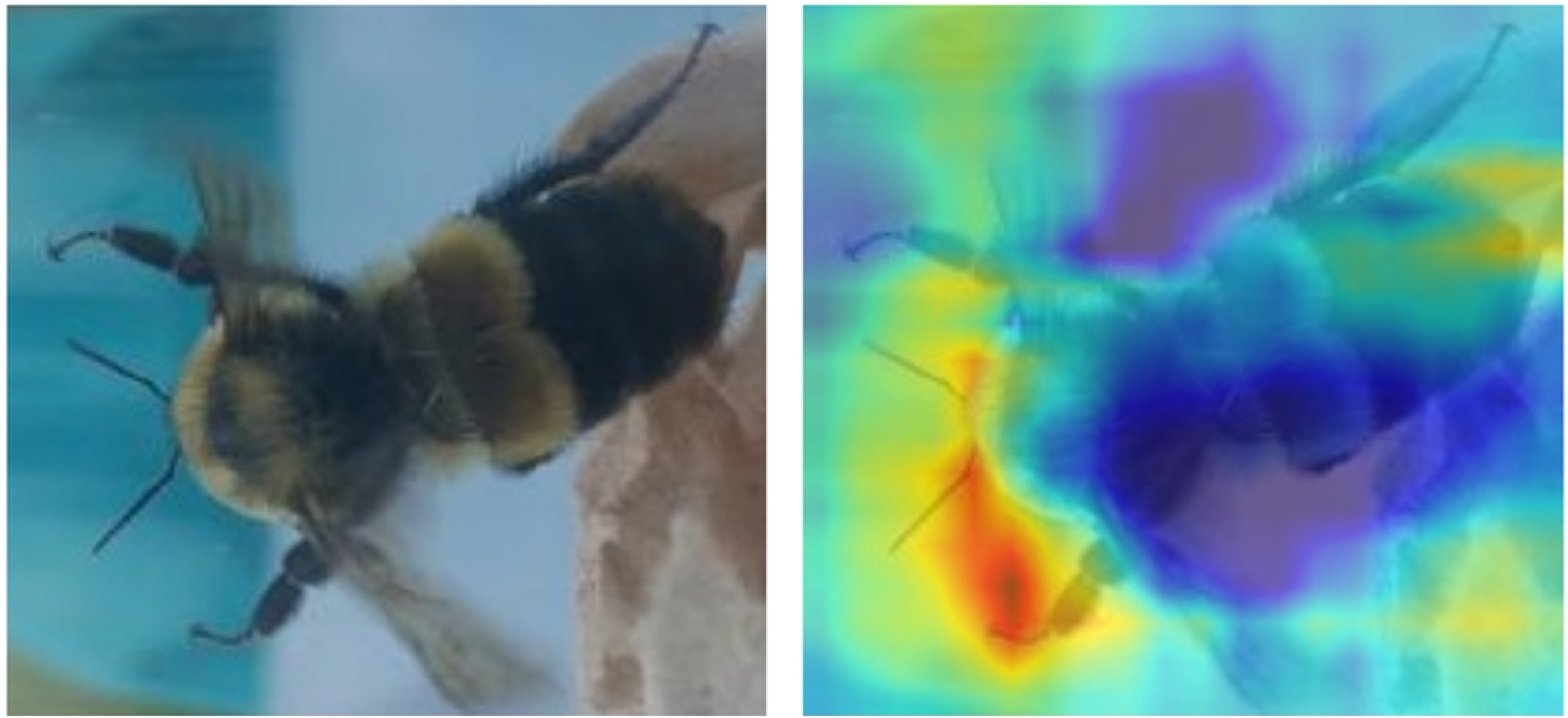
Grad-CAM analysis for EfficientNetV2M on two input images.

## 1 Conclusion

Our research addresses the critical challenges in automated bumblebee species identification by proposing an enhanced computer vision pipeline. By integrating object detection with segmentation, the pipeline effectively mitigates issues related to noisy backgrounds, poor resolution, and image quality related bottlenecks in existing methodologies. The classification model, capable of identifying the top *k* species, demonstrates robust performance across complex image datasets. In cases where the baseline approach outperforms, the use of a state-of-the-art explainable AI model provides valuable insights into the discrepancies. The results highlight the potential of the proposed approach to advance automated species identification, particularly in biological applications requiring precise and reliable classification of multiple objects in challenging imaging conditions.

